# Raman spectroscopy reveals growth phase-dependent molecular differences in bacterial membrane vesicles

**DOI:** 10.1101/2025.07.04.663173

**Authors:** Lennart Christe, Annika Haessler, Stefanie Gier, Bernd Schmeck, Nathalie Jung, Maike Windbergs

## Abstract

Bacterial membrane vesicles (BMVs) have attracted significant attention as highly efficient transport vehicles for molecules crossing biological barriers and as key mediators in infection processes. As interest in BMVs increases, the need for standardized isolation protocols and comprehensive analytical approaches becomes apparent. This study introduces Raman spectroscopy as a novel, chemically selective monitoring approach for the analysis of subtle biochemical changes in BMVs across different bacterial growth phases. BMVs derived from *Pseudomonas aeruginosa*, a Gram-negative human pathogen responsible for severe nosocomial infections, were isolated at six different time points and analyzed via established physicochemical and functional assays, as well as Raman spectroscopy. While established analytics revealed growth phase-dependent variations in protein content, surface charge, and immunogenic effects on human immune cells, Raman spectroscopy enabled the comprehensive analysis of molecular-level changes between isolation time points. Significant shifts in protein-to-lipid ratios, higher lipid saturation, and changes in protein secondary structure were detected in BMVs isolated from later growth phases. Further, the absence of spectral markers for nucleic acids enabled the identification of BMVs as outer membrane vesicles. These findings emphasize the critical influence of the isolation time point on BMV properties and highlight Raman spectroscopy as a powerful tool for semi-quantitative chemical profiling, revealing minuscule yet biologically significant changes in BMVs depending on isolation time points.

## Introduction

Vesicular transport systems are indispensable to all forms of life, fulfilling critical tasks in cellular communication and environmental interactions (1). Vesicles typically range from 20 to 200 nm in size and consist of a lipid bilayer, effectively shielding the enclosed molecules from external influences (2). This allows even fragile and enzymatic degradation-prone molecules, such as nucleic acids and proteins, to be transported (2,3). Additionally, vesicles can bypass cellular membranes and even cross biological barriers, such as endothelia and epithelia, making it possible for non-membrane-permeable substances to traverse these otherwise impermeable boundaries. Membrane proteins play a pivotal role in this process, facilitating receptor-mediated endocytosis (4,5). These extraordinary features of vesicles make them an attractive focus for research, potentially serving as a natural model for the development of biomimetic drug delivery systems, which are already being explored and applied (3, 6).

For prokaryotes, vesicles are vital not only for nutrient uptake and signal transduction but also for different defense mechanisms (2). Hence, virulence factors or antibiotic resistance-mediating enzymes are packaged within bacterial membrane vesicles (BMVs), which bud off the bacterial membrane (8–10). The Gram-negative bacterium *Pseudomonas aeruginosa* is particularly adept at transferring virulence factors during infection using BMVs, often causing severe pneumonia (9,11–14). Its high pathogenicity, combined with resistance to various antibiotics, classifies it as one of the highly problematic ESKAPE pathogens, together with *Enterococcus faecium, Staphylococcus aureus, Klebsiella pneumoniae, Acinetobacter baumannii*, and *Enterobacter* species (12,15). Therefore, vesicles originating from *P. aeruginosa* are particularly intriguing for further investigation.

As a Gram-negative bacterium, *P. aeruginosa* is enveloped by an inner and outer cell membrane separated by a relatively thin peptidoglycan cell wall, and its secreted vesicles are classified according to their respective budding origin (16). Vesicles formed through outward bulging of the outer membrane are referred to as outer membrane vesicles (OMVs), while those involving both membranes are termed outer-inner membrane vesicles (OIMVs) (11,17,18). Additionally, vesicular bodies can be formed by the lytic breakdown of cells (19,20). A detailed classification of bacterial vesicles has been described in literature, but for the sake of simplicity, all types will be referred to as bacterial membrane vesicles (BMVs) throughout this text (2). Not only the vesicle biogenesis but also the physiological state of the bacterium itself impact the chemical composition of the vesicular structure and its cargo. Bacterial cells typically undergo four consecutive growth phases: a brief lag phase, during which they adapt to their environment; an exponential log phase, characterized by rapid growth; a stationary phase, where population growth is balanced by cell death due to spatial or nutrient limitations; and finally, a death phase, when nutrient depletion or the buildup of toxic waste products causes a decline in the population (21–23). In many publications focusing on BMVs, the influence of these growth phases has received little attention, resulting in studies not specifying the exact time of isolation, consequently making it challenging to compare and reproduce BMV characteristics. Analytical challenges further impede the comparison: while standard approaches focus on fast, user-friendly physicochemical characterizations that omit discriminative chemical features of isolates, more advanced approaches like liquid chromatography-mass spectrometry (LC-MS) allow for in-depth characterization at the cost of labor-intensive sample preparation and destruction of the sample (24–27). Thus, new approaches for BMV characterization, such as Raman spectroscopy, which balances accessible experimental implementations with chemically discriminative analyses, are needed. This spectroscopic technique is based on the Raman effect, where light is partially inelastically scattered when interacting with matter. The resulting wavelengths of the scattered photons depend on energy changes in excited molecules, providing information about the chemical composition and molecular constitution (28). Raman spectroscopy, thereby, does not require extensive sample preparation, providing a label-free non-destructive analysis approach (29–32). Despite its potential, Raman spectroscopy has only been applied sparingly in the literature for the characterization of BMVs (33,34).

In order to shed light on the question of how the isolation time point alters BMV characteristics and how these changes can be revealed, this study examined BMV isolates of *P. aeruginosa* in planktonic culture at six different isolation time points throughout 24 hours. Isolates were analyzed using established physicochemical characterization techniques such as Dynamic Light Scattering (DLS) and Atomic Force Microscopy (AFM), as well as functional characterization by assessing the immunogenic potential using human macrophages. These analyses were complemented with Raman spectroscopy to reveal subtle chemical differences in BMV isolates and to offer a more comprehensive understanding of the molecular changes in BMVs depending on the isolation time point.

## Materials & Methods

Unless otherwise stated, materials were obtained from Sarstedt (Nümbrecht, Germany). Ultrapure water was provided by a PURELAB 2 Flex water purification system (Veolia Water Technologies GmbH, Celle, Germany).

### Bacteria cultivation

*Pseudomonas aeruginosa* ATCC 27853 (Thermo Scientific, Waltham, USA) was streaked onto Nutrient Broth (NB) agar plates made (Oxoid Ltd., Basingstoke, UK) and incubated overnight at 37 °C. Pre-cultures were prepared by inoculating 20 mL of Difco Nutrient Broth medium (Becton, Dickinson and Company, Sparks, USA) with a single colony and growing them for 16 h at 37 °C and 180 rpm. Main cultures (500 mL) were adjusted to a starting optical density (OD_600_) of 0.02 and cultivated at 37 °C and 180 rpm for 4, 8, 12, 16, 20, and 24 h, respectively. Bacterial growth was tracked by OD_600_ measurements every 2 h.

### Sulpho-phospho-vanillin assay

As a calibration reference, 1,2-dioleoyl-sn-glycero-3-phosphocholine (Lipoid GmbH, Ludwigshafen, Germany) liposomes were manufactured by lipid film hydration and prepared in concentrations ranging from 0.1 to 1 mg/mL. 50 µL of samples and calibration standards were mixed with 250 µL of 96% sulfuric acid (Merck KGaA, Darmstadt, Germany) and heated at 90 °C for 20 min. In a 96-well plate, 110 µL of 0.2 mg/mL vanillin in 17% phosphoric acid (both from Merck KGaA, Darmstadt, Germany) were added to 220 µL of the heated mixture and incubated at room temperature for 10 min. The plate was afterwards measured at 540 nm in a Tecan Spark multimode plate reader (Tecan Group AG, Männedort, Switzerland).

### Isolation of bacterial membrane vesicles

Bacterial cultures were first centrifuged at 15,000 x g for 10 min at 4 °C in a tabletop centrifuge (Heraeus Megafuge 16R, Thermo Scientific, Waltham, USA) to remove bacterial mass. The supernatants were filtered through 0.22 µm PES bottle top filters (Fisher Brand, New Hampshire, USA) and concentrated approximately 10-fold by tangential flow filtration (VivaFlow 200, 100 kDa cutoff, polyethersulfone, Sartorius, Göttingen, Germany). The concentrate was then ultracentrifuged for 2 h at 150,000 *xg* and 4 °C in a Beckman Optima LE-80K (Beckman Coulter, Brea, USA). The supernatant was discarded, and the pellet was washed twice in 1 mL ultrapure water. Afterward, the BMV pellet was resuspended in 1 mL ultrapure water and stored at 4 °C for a maximum of 1 week.

### Protein and LPS quantification

Protein quantification was conducted with the Pierce BCA Protein Assay Kit (Thermo Scientific, Waltham, USA), following the provided manufacturer’s instructions. BMV isolates were either untreated or supplemented with 2% w/w sodium dodecyl sulfate (Merck KGaA, Darmstadt, Germany). Accordingly, the Pierce Chromogenic Endotoxin Quant Kit (Thermo Scientific, Waltham, USA) was employed to identify lipopolysaccharide concentrations. The manufacturer’s instructions were modified by using lower volumes (80%) while maintaining the same ratios. Samples were diluted to a protein concentration of 10 µg/mL, as determined previously by the BCA method, ensuring that all samples fell within the linear range of the LPS test kit.

### Dynamic light scattering and electrophoretic light scattering

The intensity-dependent average hydrodynamic diameter, polydispersity index (PDI), and zeta potential were determined using a Zetasizer NanoZS system (Malvern Pananalytical, Malvern, USA). Samples yielded directly after BMV isolation were diluted in a ratio of 1:20 in ultrapure water and measured at 25 °C in polystyrol cuvettes.

### Atomic force microscopy

Mica substrates were prepared by attaching Muscovite Mica (12 mm diameter, 0.15-0.21 mm thickness, Electrion Microscopy Sciences, Hatfield, USA) onto microscope slides (adhesive silanized, 75×25×1 mm) followed by rinsing the mica with ethanol 96% v/v (VWR International, Darmstadt, Germany). The mica was freshly cleaved by rapidly removing a tape strip from the surface. Before placing samples on the substrate, their protein content was adjusted to 0.1 µg/mL (previously determined by BCA assay). Then, 20 µL were added to the mica surface and allowed to air dry. Before imaging, the loaded slides were stored in a desiccator overnight. All preparation steps were performed under a laminar airflow bench to exclude contaminant deposition. AFM measurements were conducted on an AFM JPK Nanowizard III (Bruker Corporation, Billerica, USA) equipped with ACTA-50 tips (tip radius < 10 nm, spring constant 37 N/m, frequency 300 kHz, 125 µm length, 30 µm width, 4 µm thickness, Al coating, APPNano, Mountain View, USA). Image settings were 2×2 µm scanning area, 1024×1024 resolution, acquired at a 0.7 Hz line rate. Collected height images were processed in Gwyddion V. 2.65 (Czech Metrology Institute among others, Jihlava, Czech Republic) by applying polynomial background removal and row alignment. Vesicles were recognized using Otsu’s automatic particle detection algorithm to calculate the artificial diameter, and subsequent total volume determination was achieved by interpolation using the Laplace function. Total volume was further divided by the number of identified vesicles and inserted into equation 1, with *V*_*L*_ being the estimated volume and the *d* the estimated diameter:

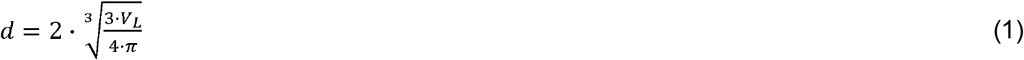

### Raman spectroscopy

BMV isolates suspended in ultrapure water were adjusted to 10 µg/mL protein, previously determined by BCA. From each BMV isolate, 10 µL were pipetted onto CaF_2_ slides (Korth Kristalle GmbH, Altenholz, Germany) and allowed to air dry. Raman spectra were collected by recording a line scan in the desired border region of the dried-drown droplet with a WITec alpha 300R+ microscope (WITec GmbH, Ulm, Germany) coupled with a 532 nm laser set to an output laser power of 10 mW before passing the objective (50x, numeric aperture 0.8, Carl Zeiss Microscopy GmbH, Jena, Germany). 10 accumulations with an integration time of 1 s were chosen for adequate signal intensities. The recorded spectra were pre-processed in the Project Four software (WiTec GmbH, Ulm, Germany) by applying background subtraction and cosmic ray removal, followed by further analysis in Matlab R2022b (Mathworks, Natick, USA) using self-written scripts. Normalization to the C-H stretching peak at 2933 cm^-1^, Savitzky-Golay filter smoothing, and spectra cropping, leaving the regions from 400 – 1800 cm^-1^ and 2800 – 3200 cm^-1^ were employed in all analysis procedures. Additional steps for 2D-COS calculation followed the procedure reported in literature (47). Raman wavenumber assignments can be found in table 1.

**Table 1.** Raman wavenumber assignments.

| Wavenumber [cm<sup>-1</sup>] | Assignment | Reference |
| --- | --- | --- |
| 748 | Tyrosine | (62) |
| 785 | Cytosine, uracil, thymine ring breathe | (33) |
| 1235 | Amide III, β sheet | (63,64) |
| 1279 | Amide III, α helix | (63,64) |
| 1302 | Lipid C-H<sub>2</sub> twisting | (65) |
| 1338 | Carbohydrate C-C | (33) |
| 1645 - 1680 | Amide I | (33,66) |
| 2850 - 2865 | Lipid C-H<sub>2</sub> | (33) |
| 2935 | Aliphatic/aromatic C-H | (33,66) |

### Differentiation of THP-1 cells to resting macrophages

THP-1 monocytes (German Collection of Microorganisms and Cell Cultures GmbH, Braunschweig, Germany) were differentiated to resting M0 macrophages by cultivating the cells with 1 mL RPMI-1640 medium + 10% fetal calf serum (FCS) supplemented with 45 ng/mL phorbol-12-myristate-13-acetate (all from Thermo Scientific, Waltham, USA). After 48 h of incubation at 37 °C and 5% CO_2_, the medium was changed to pure RPMI/FCS, and the cells were further incubated for 24 h. Differentiation to resting M0 macrophages was microscopically validated.

### Evaluation of cytotoxicity

Cytotoxicity of BMVs was analyzed using the 3-(4,5-dimethylthiazol-2-yl)-2,5-diphenyltetrazolium bromide (MTT) assay. THP-1 cells were seeded in a 96-well plate at a density of 40,000 cells per well and differentiated as described above. Subsequently, the resulting M0 macrophages were incubated with BMVs isolated after 4, 8, 12, 16, 20, and 24 h at concentrations ranging from 0.05, 0.5, and 5 µg/mL for 24 h (37 °C and 5% CO2). Afterwards, cells were washed with 1x PBS, and MTT reagent (final concentration 1 mg/ml in RPMI) was added to each well. After incubation for 4 h (37 °C, 5% CO_2_), the supernatant was discarded, and the formazan crystals were dissolved in dimethyl sulfoxide (Merck KGaA, Darmstadt, Germany). Absorbance was measured at 570 nm using the Spark multimode microplate reader (Tecan, Männerdorf, Switzerland). Cells treated with 1% TritonX100 (v/v in RPMI) or untreated cells served as positive and negative controls, respectively, and the resulting cell viability was calculated according to equation 2.

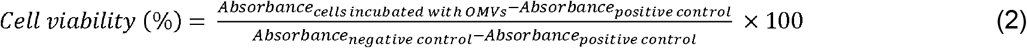

### Enzyme-linked immunosorbent assays (ELISA) of cytokines

The immunogenic potential of BMVs was evaluated via ELISA. Therefore, THP-1 cells were seeded at 350,000 cells per well in a 12-well plate and differentiated as described above. The resulting M0 macrophages were treated with BMVs (1 µg/mL protein concentration) for 6 h. Supernatants were subsequently collected, centrifuged (10.000 x g, 10 min, 4 °C), and stored at -80 °C until further analysis. Cytokine quantification was performed using the uncoated ELISA kits for IL-1β, IL-6, and TNFα following the manufacturer’s instructions (Invitrogen, Waltham, USA).

### Statistical analysis

Statistical evaluation of data was performed with GraphPad Prism 10.2.3 (GraphPad Software, Boston, USA). Ordinary one-way ANOVA coupled with Tukey’s multiple comparisons test was applied to neighboring data to test for statistical significance, highlighting changes in small time intervals. For the ELISA results, comparisons between all data points were exceptionally performed to underscore holistic trends in cytokine results. Results are shown as mean values, and error bars indicate standard deviation. Levels of statistical significance are displayed as ns (not significant), *(p < 0.05), **(p < 0.01), *** (p < 0.001) or **** (p < 0.0001).

## Results

*P. aeruginosa* was grown in nutrient broth medium over 24 h with BMVs harvested every 4 h, and bacterial growth was tracked by measuring the optical density at 600 nm (OD_600_) in 2 h intervals (fig. 1A). After a short lag phase, which lasted approximately 2 h, exponential growth was observed until 12 h of total incubation time, whereafter the OD_600_ remained consistent, indicating the stationary phase. BMV yields were quantified by determining protein amounts using a bicinchoninic acid (BCA) assay (fig. 1B). Following the logarithmic characteristics of the growth curve, protein concentrations in BMV isolates increased exponentially. Although adding 2% w/w sodium dodecyl sulfate (SDS) was reported to restrain interferences between non-protein compounds and the assay, no differences were observed in this study (35). Furthermore, lipopolysaccharide (LPS) content was assessed, revealing a slight decrease in the LPS amount over the course of bacterial growth (fig. 1C).

**Fig. 1.**
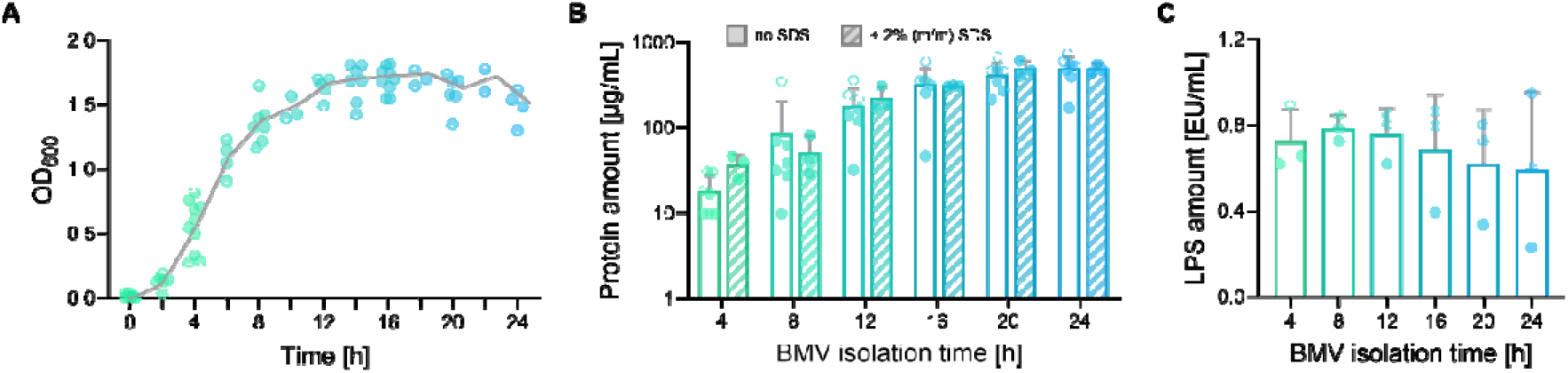
A. Evaluation of *P. aeruginosa* bacterial growth by OD_600_ measurements from 0 - 24 h. The depicted line follows mean values for each time point (n = 5). **B**. Protein quantification of BMV samples for the isolation time points 4, 8, 12, 16, 20, and 24 h. Empty bars display protein levels in untreated samples, whereas hatched bars show concentrations in samples pre-treated with 2% w/w SDS (no SDS: n = 6, with SDS: n = 3). **C**. LPS quantification of BMV samples for the isolation time points 4, 8, 12, 16, 20, and 24 h (n = 3). Error bars in all graphs indicate standard deviation. Statistical analysis of B-C in tab. S1.

Atomic force microscopy (AFM) was employed to evaluate the dimensional properties of the isolated BMVs. Exemplary records are shown in fig. 2A for all isolation times. AFM confirmed the presence of spherical objects and the absence of any unwanted bacterial remains such as pili or flagella. The average height of the dehydrated vesicles is depicted in fig. 2B. Height values were consistent at around 6 nm for all time points. Based on these measurements, diameter values were calculated from the volume covered by the deposition area, assuming the vesicles to be perfect spheres before deposition (fig. 2C). For all isolation times, comparable artificial diameters were obtained, ranging from 9 nm to 14 nm. These analyses were complemented by dynamic light scattering, which determined hydrodynamic diameters between 30 and 40 nm for all BMV isolates (fig. 2D). Using DLS measurements, the increased polydispersity of BMV populations was detected in dependence on the bacterial growth phase, with polydispersity index (PDI) values rising from 0.2 to 0.5. However, it could be noted that fluctuations in recorded PDIs decreased with time (fig. 2E). Assessing the zeta potential with electrophoretic light scattering (ELS) revealed that surface charges of BMVs increased throughout bacterial growth, starting at -22 mV and reaching values up to approximately -40 mV (fig. 2F).

**Fig. 2.**
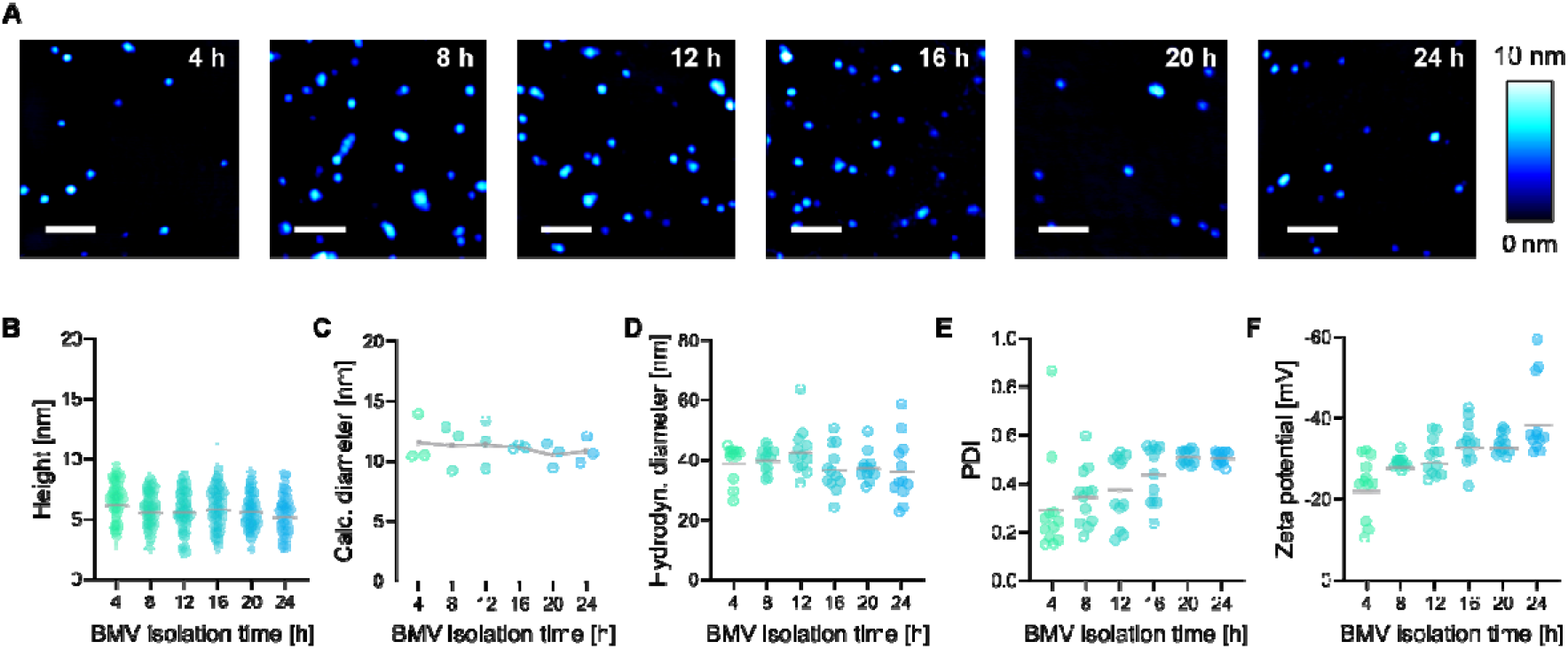
A. Representative AFM height images of dehydrated BMVs from all isolation time points (4, 8, 12, 16, 20, and 24 h). Color bar on the right indicates height-color-correlation, scale bars indicate 200 nm (total image size 1×1 µm). **B**. Height values obtained from AFM images (n = 3, 30 particles measured per n). **C**. Calculated theoretical diameter from estimated volumes covered by dehydrated BMVs (n = 3). **D**. Hydrodynamic diameter determined by DLS (n = 4, each performed in triplicate). **E**. Polydispersity index determined by DLS. Gray lines represent the mean values at the respective isolation times (n = 4, each performed in triplicate). **F**. Zeta potential determined by ELS (n = 4, each performed in triplicate). Gray lines indicate the mean values at the respective isolation times. Statistical analysis of B-F in tab. S2.

For Raman spectroscopy, all BMV suspensions were dried on calcium fluoride (CaF_2_) slides, creating a deposition of BMV material according to the coffee ring effect (fig. 3A). Systematic analysis of the border region of the dried sample was performed to determine areas containing BMVs for the following measurements. Raman spectra were recorded in parallel lines at increasing distances from the border. The selected six representative lines were distributed over the border region based on visible alterations, allowing reproducible measurements of subsequent samples. As demonstrated in fig. 3B, the respective border regions not only appeared different in bright field images but also exhibited distinct Raman spectra, indicating variations in their chemical composition. The mean spectra of line scans 1 and 2 shared identical spectral features, although both lacked protein-associated peaks, such as the prominent amide I band at 1650-1680 cm^-1^, suggesting the absence of BMV structures in these areas. Moving to more inner border districts, spectra from lines 3-5 indicated BMV deposition, as they contained a significant amide I signal; however, they exhibited variations in diverse peaks, such as the lipid C-H peak at 2850 cm^-1^ and the tyrosine peak at 748 cm^-1^. Spectra collected on line 6 at the inner border edge gave weak signals, resulting in a comparably high standard deviation and indicating low sample density. Among all detected peaks, some match with those outstanding in lines 1 and 2 (e.g., at 854 cm^-1^, 939 cm^-1,^ and 1046 cm^-1^) without appearing in lines 3-5.

**Fig. 3.**
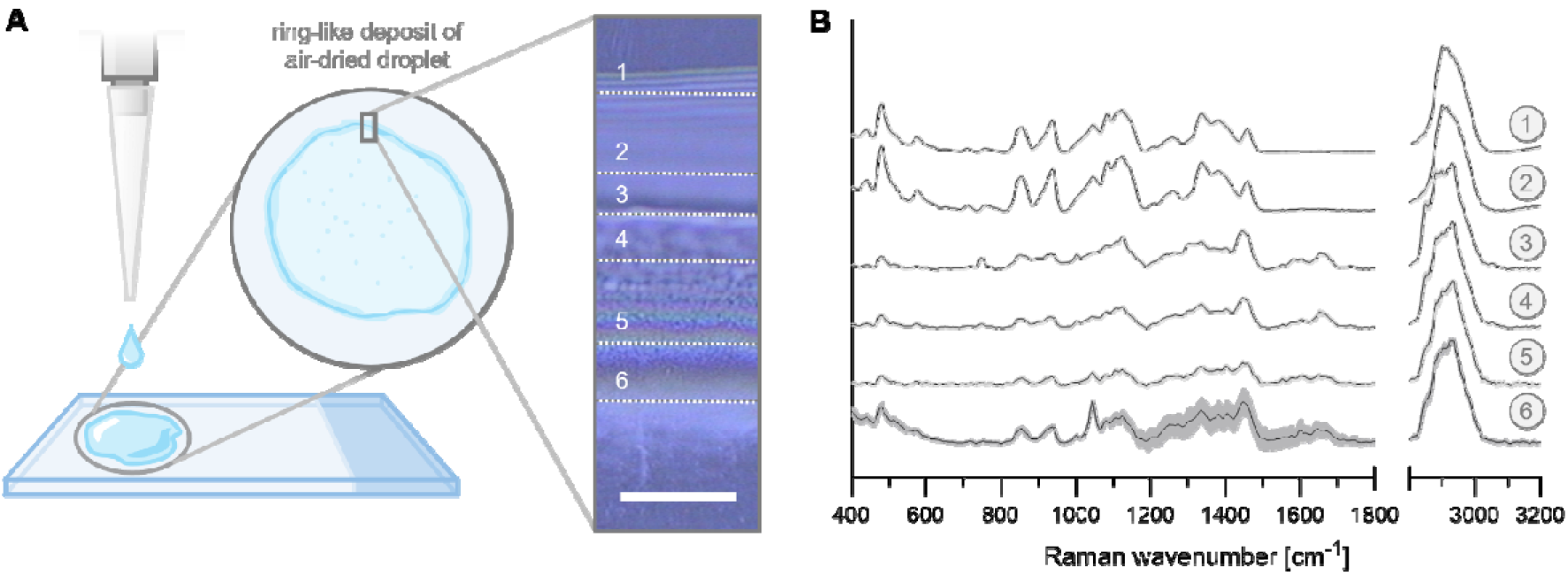
A. Exemplary border analysis of the coffee ring formed out of dried-down BMV suspension (isolated after 20 h of bacterial growth). Schematic illustration of deposition with a bright field image showing the border of the dried droplet (scale bar 20 µm, outer to inner border from top to bottom), wherein six representative lines for Raman analysis are marked. **B**. Mean Raman spectra from the six line scans are displayed with their respective standard deviation and numeration corresponding to the respective lines in the bright field image.

Based on these results, all further scans were performed as line scans in the exact middle of the visually highlighted structured area (between lines 3 and 5), ensuring comparability and uniformity for all data.

Raman spectra of all BMV isolates were collected, and the resulting mean spectra are depicted in fig. 4A, showing annotations for all relevant peaks. By performing a principal component analysis (PCA), the clustering of the six different BMV isolations was illustrated (fig. 4B). The distribution reveals variations, especially at earlier isolation time points (4 h, 8 h), with the most significant separation along PC1, accounting for 56.4% of total variance. Apparent clustering was observed at especially early time points, whereas overlapping increased for later sampling points (16 h and 24 h). Group-internal deviations decreased with longer growth duration; however, distinguishing between later isolates became increasingly more complex.

**Fig. 4.**
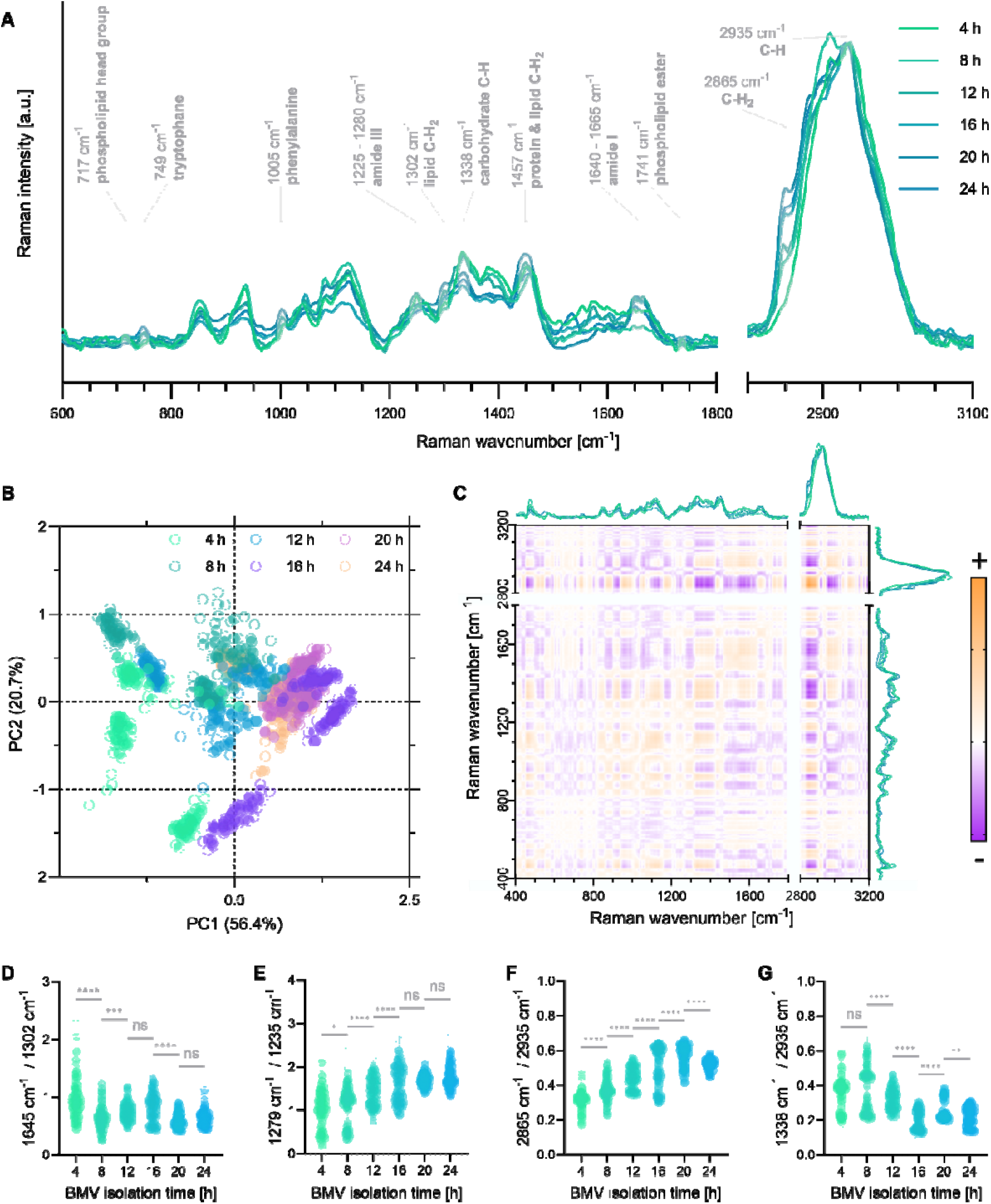
A. Mean Raman spectra of all BMV isolates normalized to the C-H peak at 2935 cm^-1^. Prominent Raman bands are annotated and marked in gray (n = 4, 50 spectra per n). **B**. Principal component analysis of all BMV spectra (n = 3, 50 spectra per n). **C**. 2D-COS analysis of mean BMV spectra of all BMV isolates. Orange: positive correlation; purple: negative correlation. **D-G**. Peak ratio analysis of distinct spectral features of BMV isolate spectra from 4, 8, 12, 16, 20, and 24 h, respectively. Statistical analysis: One-way ANOVA and Tukey’s post-hoc test (n = 4, full statistical analysis in tab. S3). Statistical significance is indicated by *(p < 0.05), **(p < 0.01), *** (p < 0.001) or **** (p < 0.0001), and ns (not significant). **D**. Protein-to-lipid ratio (amide I band at 1645 cm^-1^, lipid C-H_2_ twisting at 1302 cm^-1^). **E**. Protein secondary structures ratio (amide III band; α-helix at 1279 cm^-1^, β- sheet at 1235 cm^-1^). **F**. Saturated lipid peak ratio (C-H_2_ bonds at 2865 cm^-1^, C-H bonds at 2935 cm^-1^). **G**. Carbohydrate peak ratio (carbohydrate C-C bonds at 1338 cm^-1^, C-H bonds at 2935 cm^-1^).

An overview of all dynamic changes in relative signal intensities between isolations was visualized using two-dimensional correlation spectroscopy (2D-COS) (fig. 4C). The symmetrical 2D-COS plot revealed all peak relationships by color-mapping positive and negative correlations of spectral information over the impact of isolation times. For example, examining the 2865 cm^-1^ (lipid C-H2 peak) from both the x-and y-plotted mean spectra reveals an intense orange blot, indicating a positive correlation. Hereby, an increase in the BMV isolation time of the regarded peak on one axis was accompanied by an increase in intensity for the desired peak on the other axis. A strong, exemplary negative correlation (purple blot) could be observed by comparing the lipid C-H_2_ intensity at 2865 cm^-1^ to the carbohydrate C-C signal at 1338 cm^-1^, where the intensity trends of both peaks are inversely correlated. The 2D visualization of time-dependent changes in the spectroscopic data set allowed the identification of relevant Raman peak pairs for subsequent peak ratio analysis. Calculating peak ratios (fig. 4D-G) enabled more precise insights into distinct spectral features and their semi-quantitative assessment throughout bacterial growth. Fig. 4D displayed the protein-to-lipid ratio by analyzing the relation of amide I intensity at 1645 cm^-1^ to lipid-assigned C-H_2_ twisting at 1302 cm^-1^, and revealed highly significant differences in protein portion based on the lipid content between isolations at 4 h and 8 h, 8 h and 12 h as well as 16 h and 20. Fig. 4E illustrates the detected shifts in predominant protein secondary structure by comparing two points in the amide III bands corresponding to α-helices at 1279 cm^-1^ and to β-sheets at 1235 cm^-1^. According to these findings, the proportion of α-helix structures in total protein structures increased with later BMV isolates, resulting in significant differences, especially between isolations up to 16 h. Furthermore, an increase in lipid saturation (fig. 4F) in vesicles throughout bacterial growth was observed when relating the C-H_2_ signal at 2865 cm^-1^ to the C-H signal at 2935 cm^-1^, revealing highly significant differences between all isolation points. Another peak ratio calculated to address the carbohydrate content in BMVs showed a decrease in sugar structures (carbohydrate C-C bonds, 1338 cm^-1^) with ongoing cultivation durations when normalized to the C-H peak at 2935 cm^-1^ (fig. 4G), which were especially prominent in later BMV isolates.

The immunogenic potential of BMVs from different isolation time points was analyzed by treating resting macrophages (M0) with 1 µg/mL BMVs for 6 h. The secretion of the pro-inflammatory cytokines IL-1β, IL-6, and TNF-α was then evaluated using enzyme-linked immunosorbent assays (ELISA). Beforehand, the cytotoxicity of BMVs was evaluated by an MTT assay, revealing no significant toxic effects on M0 macrophages at the chosen concentration (see fig. S1). BMVs from different isolation time points influenced the respective cytokine releases differently, depending on the investigated cytokine. IL-1β levels consistently decreased when M0 macrophages were treated with later BMV isolates, reaching cytokine concentrations that do not differ significantly from the negative control in BMVs from 20 h and 24 h. The IL-1β release of nearly 45 pg/mL for BMVs isolated at 4 h exceeded the immune response observed in the positive control by almost 80%. In contrast, BMVs from every isolation triggered comparable IL-6 secretion, inducing the release of over 100 pg/mL. Regarding the release of TNF-α, all isolations induced a significant immune response, and a slight increase was detected when comparing the 4 h and the 24 h isolate, where TNFα levels rose from approximately 1800 pg/mL to 2300 pg/mL, resembling an increase of about 28%.

## Discussion

Understanding the dynamics of membrane vesicle release throughout the lifecycle of *P. aeruginosa* is crucial to deciphering its adaptive strategies. Here, we employed a combined approach utilizing established biochemical assays and, for the first time in this context, Raman spectroscopy to elucidate vesicle characteristics at defined time points. By tracking the optical density at 600 nm throughout 24 h, the temporal sequence of different growth phases was recorded (fig. 1A). The exponential character of the growth curve corresponded to the trend in quantified protein concentrations in all collected vesicle isolates (fig. 1B), revealing a correlation of total bacteria mass to isolated bacterial vesicle amounts. In addition to protein quantification, lipopolysaccharide contents were evaluated, showing an exponential trend with a slight non-significant decrease at later isolation times (fig. 1C).

After BMV isolation, the purity of isolates was assessed using DLS and AFM analysis. Both measurements highlighted the absence of any larger biological debris, such as bacteria pili or flagella (fig. 2). The average hydrodynamic diameter was approximately 40 nm, remaining almost consistent over the course of growth. Regarding the polydispersity index, variations in BMV size increase with later isolation times, revealing a broad size distribution for BMVs. According to the literature, bacterial vesicles from *P. aeruginosa* range in hydrodynamic diameter from 25 to 200 nm, in compliance with the measured sizes in this study (36–38). AFM measurements exhibited height values of BMVs of around 6 nm, which were drastically lower than previously reported values, stating heights for OMVs from *P. aeruginosa* of 10 – 30 nm (38). However, these height values were detected when the vesicles were retained in a hydrated state, whereas in this study, the vesicles were dried after deposition. The hereby acquired values can be attributed to twice the thickness of a lipid bilayer (approximately 4 nm), further flattened by deformation processes during drying and storage (39,40). To obtain a more meaningful indicator of the size changes across isolation time points from AFM data, an artificial diameter was calculated from the volume covered by the deposition area, assuming the vesicles to be perfect spheres (fig. 2C). This attempt has previously been proven to comply with particle sizes acquired by cryogenic electron microscopy, although particles need to be hydrated for best accuracy (40). In this case, the calculated diameters rather provide a means for relative comparison rather than absolute determination. Regardless of the employed method, no significant changes in BMV size were observed between the different isolation time points.

For Raman spectroscopic evaluations, BMV isolates were drop-casted and fully dried before analysis. During the drying process, particle-like structures accumulate in the border region of dried droplets, a phenomenon known as the coffee ring effect (41,42). Raman spectra were therefore recorded in this region, allowing for the best possible comparison between different samples. Analysis of the border region, however, revealed notable differences in the acquired Raman spectra depending on the position of spectra acquisition (fig. 3). In the outer regions 1 and 2, similar spectra were recorded that can be assigned to carbohydrate structures consisting of 1,4-linked α-glucose polymers like glycogen or amylose (43,44). Although the secretion of such polysaccharides from *P. aeruginosa* has not yet been reported, extracellular glycogen accumulation during growth in rich medium conditions was already observed for its direct relative, *P. fluorescens* (45). A deposition overlap of BMV structures with such carbohydrates cannot be excluded; nonetheless, spectra recorded across the border area revealed distinct signals of both entities in the analyzed regions. Summarizing these findings, the center area of the border is the most suitable for collecting comparable spectra of deposited bacterial vesicles, as it offers the densest deposition layer, preventing susceptibility to variations in composition. Still, exact spatial discrimination of coffee ring border entities remains a major challenge with potential implications for the gathered results. Further purification of isolated BMVs could also reduce confounding interferences, as spectral signatures not associated with bacterial vesicles were detected.

Regarding the Raman spectra of all BMV isolates (fig. 4A), the absence of nucleic acid-associated peaks, such as cytosine, uracil, and thymine ring breathing modes at 785 cm^−1^, indicates that DNA and RNA were not present in the analyzed vesicles (33). This suggests that lysis-derived vesicles, typically carriers of nucleic acids, are present only in minimal amounts, and the majority of isolated vesicles are outer membrane vesicles, which lack genetic material. By monitoring the presence of nucleic acids, Raman spectroscopy can be employed to discriminate between lytic vesicles and OMVs as an alternative to PAGE analysis for confirming vesicle populations after fractionation (46). Principal component analysis (PCA) was applied to assess the consistency and variance across biological replicates. Early isolation time points exhibited higher variability (fig. 4B), likely due to lower vesicle yields during the early growth phases when protein concentrations are still increasing (fig. 1A, B). Additionally, the rapid metabolic and structural changes in bacteria during early exponential growth may cause subtle timing differences in sampling, which can disproportionately affect vesicle composition. PCA also enabled discrimination between vesicle populations from different isolation time points. Since PC1 captured the majority of variance in the dataset, its loading spectra were further analyzed to identify the primary contributors to this separation (fig. S3). The most prominent contribution was observed at 2850 cm^−1^, corresponding to symmetric CH_2_ stretching vibrations, a spectral marker for lipids. This indicates that lipid content is a key factor differentiating BMVs across isolation times and is a major driver of spectral variance along PC1. To further explore correlations between spectral features across the isolation timeline, symmetrical two-dimensional (2D) correlation spectroscopy was employed. This method highlights dynamic relationships between Raman peaks in response to systematic changes, in this case, bacterial growth duration prior to vesicle isolation (47). Peak ratios providing crucial discriminative information about chemical composition between BMV isolates were identified. Viewing the protein-to-lipid ratio in fig. 4D, protein content decreased relative to the lipid amounts. Unfortunately, no suitable biochemical assays are available for conducting lipid quantifications in such highly complex samples that could verify a similar trend with established assays. Both the Stewart and the sulpho-phospho-vanillin (SPV) assays, two established and commonly applied methods for quantifying phospholipids in microorganisms or liposomes, were highly prone to harsh deviations and interfering substances, thus yielding unreliable results for bacterial membrane vesicles (the Stewart assay failed; SPV assay results are shown in fig. S2) (48,49). However, protein amounts are easily accessible by employing the widely accepted BCA assay, a state-of-the-art method for assessing bacterial vesicle yields (fig. 1B) (13,50). Combining these results with Raman spectroscopy circumvents the limitations of current lipid assays and enables conclusions to be drawn regarding lipid content. Diving deeper into protein analysis by evaluating the amide III signals, α-helical structures were found to be more prominent compared to β-sheets with continuous bacterial growth (fig. 4E). Typical transmembrane proteins present in bacterial vesicles, such as porins, primarily consist of β-barrel conformations in their transmembrane domain (9,51). This reveals higher α-helix proportions combined with overall decreasing protein amounts in later vesicle isolates, potentially indicating a decrease in transmembrane protein abundance. Furthermore, viewing the saturation status of lipids in BMVs (fig. 4F), later vesicle isolates show higher lipid saturation than earlier ones. It has been reported that the remodeling of lipid bilayer structures during bacterial growth is conducted in response to environmental stimuli (52). Thereby, increased phospholipid integration into the outer membrane, as well as hepta-acylation of lipid A in LPS molecules, modifies membrane stability and leads to higher abundances of saturated fatty acids in subsequently released BMVs (53,54). The saturation deviations between the isolation time points could thus be attributed to such changes in the bacteria’s lipid bilayer composition.

Lower carbohydrate levels were detected in samples from later isolation times (fig. 4G), corresponding to the previous LPS quantification (fig. 1C). As LPS molecules are typically classified as pro-inflammatory stimuli, it is notable that their decreasing levels did not uniformly lead to declining secretions of the analyzed classical pro-inflammatory cytokines, TNF-α, IL-1β and IL-6, by resting macrophages treated with the six BMV isolates (see fig. 5) (55–57). *P. aeruginosa* OMVs are reported to activate the Toll-like receptor 4 (TLR-4) pathway in eukaryotic cells, causing secretion of pro-inflammatory cytokines such as IL-1β and IL-6, which was also observed in this study (58). Interestingly, there is no apparent trend in triggering a general immune response; instead, each observed cytokine exhibits a distinct course of release. Considering the release kinetics of the cytokines, TNF-α is a fast-acting mediator that is immediately formed when an immune response is triggered. In contrast, IL-1β levels rise more slowly and are sustained for a more extended period, typically around 24 hours. IL-6 release lies midway between the two (59). Interestingly, ELISA data revealed a gradual decrease in secreted 1L-1β after treatment of resting macrophages with BMVs collected at later time points, whereas Il-6 and TNF-α secretion remained consistently high for all samples. The results could indicate a slower or delayed immune response in BMVs sampled at later time points, potentially due to different compositions of the vesicles triggering distinct inflammatory pathways. Although further analyses are required to reveal the exact underlying mechanisms, these findings further emphasize the highly variable characteristics of BMVs collected after different bacterial growth times.

**Fig. 5.**
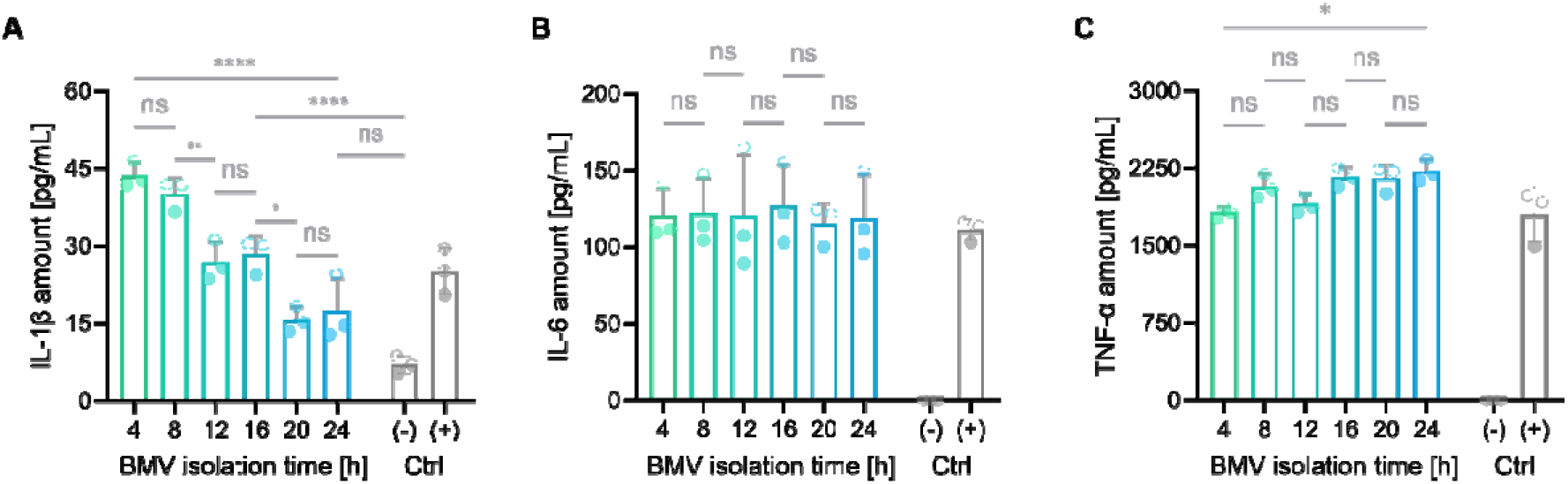
Enzyme-linked immunosorbent assays of M0 macrophages treated with 1 µg/mL of the respective BMV isolate (4, 8, 12, 16, 20, and 24 h). Investigated cytokines are IL-1β (**A**), IL-6 (**B**), and TNF-α (**C**). Negative (Ctrl-, vehicle) and positive control (Ctrl+, 50 ng/mL LPS) are depicted in gray bars. Error bars indicate standard deviation. Statistical analysis: One-way ANOVA and Tukey’s post-hoc test (n = 3, full statistical analysis in tab. S4). Statistical significance is indicated for selected comparisons by *(p < 0.05), **(p < 0.01), *** (p < 0.001) or **** (p < 0.0001), and ns (not significant),.

In our study, employing the applied analytical toolset enabled precise and specific discrimination of bacterial membrane vesicles secreted during the different growth stages of *P. aeruginosa*. In addition to increasing lipid levels, diminished carbohydrate and protein amounts were detected, while α-helical protein structures became simultaneously more prominent. Additionally, immune responses observed in resting macrophages triggered by BMVs differ significantly between vesicles harvested at various growth stages. These findings highlight the importance of uniform and standardized isolation procedures when working with bacterial vesicles. As even minor deviations in the isolation time point were revealed to have a highly significant impact on the vesicles’ attributes, experimental statements might be dramatically influenced. Raman spectroscopy has been demonstrated to be a readily accessible technique that offers valuable insights into various molecular aspects of BMVs, including lipid saturation and protein conformation, which would otherwise be limited to LC-MS applications. These features qualify Raman spectroscopy as a powerful tool for the quality control of vesicular systems, with already reported applications for liposomes intended for therapeutic use or the classification of exosomes (60, 61). There are notable drawbacks, though, as the exact quantification of particular contents is hardly possible, and the direct identification of compounds, such as specific proteins, cannot be achieved. However, when complemented with physicochemical characterization in the form of DLS, ELS, and AFM, a comprehensive evaluation of bacterial vesicles can be achieved with minimal sample volume requirements. Bacterial vesicles will remain a hot scientific topic for the near future, and with growing interest, ensuring comparability by defining strict standards regarding experimental protocols and applying high-fidelity analytics will be key to bundled and targeted scientific efforts.

## Author Contributions

**Lennart Christe**: Conceptualization, Data curation, Formal Analysis, Investigation, Methodology, Software, Validation, Visualization, Writing - original draft, Writing - review & editing. **Annika Haessler**: Formal Analysis, Investigation, Validation, Visualization. **Stefanie Gier:** Supervision, Writing – Review & Editing. **Bernd Schmeck:** Validation, Writing – Review & Editing. **Nathalie Jung**: Conceptualization, Investigation, Supervision, Writing - original draft, Writing - Review & Editing. **Maike Windbergs**: Conceptualization, Funding acquisition, Project administration, Resources, Writing - Review & Editing.

## Literature

1. Van Niel G, D’Angelo G, Raposo G. 2018. Shedding light on the cell biology of extracellular vesicles. Nat Rev Mol Cell Biol. 19(4):213–28.

2. Toyofuku M, Schild S, Kaparakis-Liaskos M, Eberl L. 2023. Composition and functions of bacterial membrane vesicles. Nat Rev Microbiol. 21(7):415–30.

3. Tenchov R, Sasso JM, Wang X, Liaw WS, Chen CA, Zhou QA. 2022. Exosomes Nature’s Lipid Nanoparticles, a Rising Star in Drug Delivery and Diagnostics. ACS Nano. 16(11):17802–46.

4. Joshi BS, de Beer MA, Giepmans BNG, Zuhorn IS. 2020. Endocytosis of Extracellular Vesicles and Release of Their Cargo from Endosomes. ACS Nano. 14(4):4444–55.

5. Karthikeyan R, Gayathri P, Gunasekaran P, Jagannadham M V., Rajendhran J. 2020. Functional analysis of membrane vesicles of Listeria monocytogenes suggests a possible role in virulence and physiological stress response. Microb Pathog. 142(104076).

6. Aytar Çelik P, Erdogan-Gover K, Barut D, Enuh BM, Amasya G, Sengel-Türk CT, et al. 2023. Bacterial Membrane Vesicles as Smart Drug Delivery and Carrier Systems: A New Nanosystems Tool for Current Anticancer and Antimicrobial Therapy. Pharmaceutics. 15(4).

7. Herrmann IK, Wood MJA, Fuhrmann G. 2021. Extracellular vesicles as a next-generation drug delivery platform. Nature Nanotechnology 2021 16:7. 16(7):748–59.

8. Cecil JD, Sirisaengtaksin N, O’Brien-Simpson NM, Krachler AM. 2019. Outer Membrane Vesicle-Host Cell Interactions. Microbiol Spectr. 7(1).

9. Zavan L, Fang H, Johnston EL, Whitchurch C, Greening DW, Hill AF, et al. 2023. The mechanism of Pseudomonas aeruginosa outer membrane vesicle biogenesis determines their protein composition. Proteomics. 23(10):2200464.

10. Schwechheimer C, Kuehn MJ. 2015. Outer-membrane vesicles from Gram-negative bacteria: Biogenesis and functions. Nat Rev Microbiol. 13(10):605–19.

11. Kadurugamuwa JL, Beveridge TJ. 1995. Virulence factors are released from Pseudomonas aeruginosa in association with membrane vesicles during normal growth and exposure to gentamicin: a novel mechanism of enzyme secretion. J Bacteriol. 177(14):3998.

12. Qin S, Xiao W, Zhou C, Pu Q, Deng X, Lan L, et al. 2022. Pseudomonas aeruginosa: pathogenesis, virulence factors, antibiotic resistance, interaction with host, technology advances and emerging therapeutics. Signal Transduct Target Ther. 7(1):1–27.

13. Augustyniak D, Olszak T, Drulis-Kawa Z. 2022. Outer Membrane Vesicles (OMVs) of Pseudomonas aeruginosa Provide Passive Resistance but Not Sensitization to LPS-Specific Phages. Viruses. 14(1).

14. Henriquez T, Falciani C. 2023. Extracellular Vesicles of Pseudomonas: Friends and Foes. Antibiotics. 12(4).

15. Pendleton JN, Gorman SP, Gilmore BF. 2013. Clinical relevance of the ESKAPE pathogens. Expert Rev Anti Infect Ther. 11(3):297–308.

16. Motta S, Vecchietti D, Martorana AM, Brunetti P, Bertoni G, Polissi A, et al. 2020. The Landscape of Pseudomonas aeruginosa Membrane-Associated Proteins. Cells. 9(11):1–17.

17. Baeza N, Delgado L, Comas J, Mercade E. 2021. Phage-Mediated Explosive Cell Lysis Induces the Formation of a Different Type of O-IMV in Shewanella vesiculosa M7T. Front Microbiol. 12:713669.

18. Roier S, Zingl FG, Cakar F, Durakovic S, Kohl P, Eichmann TO, et al. 2016. A novel mechanism for the biogenesis of outer membrane vesicles in Gram-negative bacteria. Nat Commun. 7:10515.

19. Toyofuku M, Cárcamo-Oyarce G, Yamamoto T, Eisenstein F, Hsiao CC, Kurosawa M, et al. 2017. Prophage-triggered membrane vesicle formation through peptidoglycan damage in Bacillus subtilis. Nat Commun. 8(1):1–10.

20. Abe K, Toyofuku M, Nomura N, Obana N. 2021. Autolysis-mediated membrane vesicle formation in Bacillus subtilis. Environ Microbiol. 23(5):2632–47.

21. Ughy B, Nagyapati S, Lajko DB, Letoha T, Prohaszka A, Deeb D, et al. 2023. Reconsidering Dogmas about the Growth of Bacterial Populations. Cells. 12(10).

22. Navarro Llorens JM, Tormo A, Martínez-García E. 2010. Stationary phase in gram-negative bacteria. FEMS Microbiol Rev. 34(4):476–95.

23. Rolfe MD, Rice CJ, Lucchini S, Pin C, Thompson A, Cameron ADS, et al. 2012. Lag Phase Is a Distinct Growth Phase That Prepares Bacteria for Exponential Growth and Involves Transient Metal Accumulation. J Bacteriol. 194(3):686.

24. Rešetar Maslov D, Rubić I, Farkaš V, Kuleš J, Beer Ljubić B, Beletić A, et al. 2024. Characterization and LC-MS/MS based proteomic analysis of extracellular vesicles separated from blood serum of healthy and dogs naturally infected by Babesia canis. A preliminary study. Vet Parasitol. 328.

25. Tréton G, Sayer C, Schürz M, Jaritsch M, Müller A, Matea CT, et al. 2023. Quantitative and functional characterisation of extracellular vesicles after passive loading with hydrophobic or cholesterol-tagged small molecules. Journal of Controlled Release. 361:694–716.

26. Tani C, Stella M, Donnarumma D, Biagini M, Parente P, Vadi A, et al. 2014. Quantification by LC-MS(E) of outer membrane vesicle proteins of the Bexsero® vaccine. Vaccine. 32(11):1273–9.

27. Wu J, An M, Zhu J, Tan Z, Y. Chen G, Stidham RW, et al. 2019. A Method for Isolation and Proteomic Analysis of Outer Membrane Vesicles from Fecal Samples by LC-MS/MS. J Proteomics Bioinform. 12(2):38.

28. Rull F. 2012. The raman effect and the vibrational dynamics of molecules and crystalline solids. European Mineralogical Union Notes in Mineralogy. 12(1):1–60.

29. Bukva M, Dobra G, Gomez-Perez J, Koos K, Harmati M, Gyukity-Sebestyen E, et al. 2021. Raman spectral signatures of serum-derived extracellular vesicle-enriched isolates may support the diagnosis of CNS tumors. Cancers (Basel). 13(6):1–19.

30. Gualerzi A, Kooijmans SAA, Niada S, Picciolini S, Brini AT, Camussi G, et al. 2019. Raman spectroscopy as a quick tool to assess purity of extracellular vesicle preparations and predict their functionality. J Extracell Vesicles. 8(1).

31. Zini J, Saari H, Ciana P, Viitala T, Lõhmus A, Saarinen J, et al. 2022. Infrared and Raman spectroscopy for purity assessment of extracellular vesicles. European Journal of Pharmaceutical Sciences. 172.

32. Liangsupree T, Multia E, Saarinen J, Ruiz-Jimenez J, Kemell M, Riekkola ML. 2022. Raman spectroscopy combined with comprehensive gas chromatography for label-free characterization of plasma-derived extracellular vesicle subpopulations. Anal Biochem. 647:114672.

33. Potter M, Hanson C, Anderson AJ, Vargis E, Britt DW. 2020. Abiotic stressors impact outer membrane vesicle composition in a beneficial rhizobacterium: Raman spectroscopy characterization. Scientific Reports 2020 10:1. 10(1):1–14.

34. Ren B, Cui L, Qin YF, Lu XY, Shi Z, Huang QS, et al. 2022. Deep Learning-Enabled Raman Spectroscopic Identification of Pathogen-Derived Extracellular Vesicles and the Biogenesis Process. Anal Chem. 94(36):12416–26.

35. Morton RE, Evans TA. 1992. Modification of the bicinchoninic acid protein assay to eliminate lipid interference in determining lipoprotein protein content. Anal Biochem. 204(2):332–4.

36. Cooke AC, Nello A V., Ernst RK, Schertzer JW. 2019. Analysis of Pseudomonas aeruginosa biofilm membrane vesicles supports multiple mechanisms of biogenesis. PLoS One. 14(2).

37. Zare Banadkoki E, Rasooli I, Ghazanfari T, Siadat SD, Shafiee Ardestani M, Owlia P. 2022. Pseudomonas aeruginosa PAO1 outer membrane vesicles-diphtheria toxoid conjugate as a vaccine candidate in a murine burn model. Sci Rep. 12(1):1–14.

38. Kikuchi Y, Obana N, Toyofuku M, Kodera N, Soma T, Ando T, et al. 2020. Diversity of physical properties of bacterial extracellular membrane vesicles revealed through atomic force microscopy phase imaging. Nanoscale. 12(14):7950–9.

39. Regan D, Williams J, Borri P, Langbein W. 2019. Lipid Bilayer Thickness Measured by Quantitative DIC Reveals Phase Transitions and Effects of Substrate Hydrophilicity. Langmuir. 35(43):13805–14.

40. Skliar M, Chernyshev VS. 2019. Imaging of extracellular vesicles by atomic force microscopy. Journal of Visualized Experiments. 2019(151).

41. Mampallil D, Eral HB. 2018. A review on suppression and utilization of the coffee-ring effect. Adv Colloid Interface Sci. 252:38–54.

42. Jeong H, Han C, Cho S, Gianchandani Y, Park J. 2018. Analysis of Extracellular Vesicles Using Coffee Ring. ACS Appl Mater Interfaces. 10(27):22877–82.

43. Kamemoto LE, Misra AK, Sharma SK, Goodman MT, Hugh LUK, Dykes AC, et al. 2010. Near-infrared micro-Raman spectroscopy for in vitro detection of cervical cancer. Appl Spectrosc. 64(3):255–61.

44. Bağcıoğlu M, Zimmermann B, Kohler A. 2015. A Multiscale Vibrational Spectroscopic Approach for Identification and Biochemical Characterization of Pollen. PLoS One. 10(9).

45. Quilès F, Polyakov P, Humbert F, Francius G. 2012. Production of extracellular glycogen by Pseudomonas fluorescens: spectroscopic evidence and conformational analysis by biomolecular recognition. Biomacromolecules. 13(7):2118–27.

46. Duong P, Chung A, Bouchareychas L, Raffai RL. 2019. Cushioned-Density Gradient Ultracentrifugation (C-DGUC) improves the isolation efficiency of extracellular vesicles. PLoS One. 14(4):e0215324.

47. Lasch P, Noda I. 2019. Two-Dimensional Correlation Spectroscopy (2D-COS) for Analysis of Spatially Resolved Vibrational Spectra. Appl Spectrosc. 73(4):359–79.

48. Di Prima G, Librizzi F, Carrotta R. 2020. Light Scattering as an Easy Tool to Measure Vesicles Weight Concentration. Membranes (Basel). 10(9):222.

49. McMahon A, Lu H, Butovich IA. 2013. The Spectrophotometric Sulfo-Phospho-Vanillin Assessment of Total Lipids in Human Meibomian Gland Secretions. Lipids. 48(5):513.

50. Bauman SJ, Kuehn MJ. 2006. Purification of outer membrane vesicles from Pseudomonas aeruginosa and their activation of an IL-8 response. Microbes Infect. 8(9–10):2400–8.

51. Chevalier S, Bouffartigues E, Bodilis J, Maillot O, Lesouhaitier O, Feuilloley MGJ, et al. 2017. Structure, function and regulation of Pseudomonas aeruginosa porins. FEMS Microbiol Rev. 41(5):698–722.

52. Juodeikis R, Carding SR. 2022. Outer Membrane Vesicles: Biogenesis, Functions, and Issues. Microbiol Mol Biol Rev. 86(4).

53. Bonnington KE, Kuehn MJ. 2017. Breaking the bilayer: OMV formation during environmental transitions. Microbial Cell. 4(2):64.

54. Giordano NP, Cian MB, Dalebroux ZD. 2020. Outer Membrane Lipid Secretion and the Innate Immune Response to Gram-Negative Bacteria. Infect Immun. 88(7).

55. Huszczynski SM, Lam JS, Khursigara CM. 2020. The Role of Pseudomonas aeruginosa Lipopolysaccharide in Bacterial Pathogenesis and Physiology. Pathogens. 9(1).

56. Meng F, Lowell CA. 1997. Lipopolysaccharide (LPS)-induced Macrophage Activation and Signal Transduction in the Absence of Src-Family Kinases Hck, Fgr, and Lyn. J Exp Med. 185(9):1661.

57. Turner MD, Nedjai B, Hurst T, Pennington DJ. 2014. Cytokines and chemokines: At the crossroads of cell signalling and inflammatory disease. BBA - Molecular Cell Research. 1843(11):2563–82.

58. Zhao K, Deng X, He C, Yue B, Wu M. 2013. Pseudomonas aeruginosa Outer Membrane Vesicles Modulate Host Immune Responses by Targeting the Toll-Like Receptor 4 Signaling Pathway. Infect Immun. 81(12):4509.

59. Medzhitov R, Janeway C. 2000. Innate immune recognition: mechanisms and pathways. Immunol Rev. 173(1):89–97.

60. Rodà F, Picciolini S, Mangolini V, Gualerzi A, Seneci P, Renda A, et al. 2023. Raman Spectroscopy Characterization of Multi-Functionalized Liposomes as Drug-Delivery Systems for Neurological Disorders. Nanomaterials. 13(4):699.

61. Parlatan U, Ozen MO, Kecoglu I, Koyuncu B, Torun H, Khalafkhany D, et al. 2023. Label-Free Identification of Exosomes using Raman Spectroscopy and Machine Learning. Small. 19(9).

62. Singh S, Verma T, Khamari B, Bulagonda EP, Nandi D, Umapathy S. 2023. Antimicrobial Resistance Studies Using Raman Spectroscopy on Clinically Relevant Bacterial Strains. Anal Chem. 95(30):11342–51.

63. Adar F, Zheng N, Wang Z, Zhao F, Li H, He H, et al. 2022. Interpretation of Raman Spectrum of Proteins. Spectroscopy. 37(2):9–13.

64. Hauptmann A, Hoelzl G, Mueller M, Bechtold-Peters K, Loerting T. 2023. Raman Marker Bands for Secondary Structure Changes of Frozen Therapeutic Monoclonal Antibody Formulations During Thawing. J Pharm Sci. 112(1):51–60.

65. Katainen E, Elomaa M, Laakkonen UM, Sippola E, Niemelä P, Suhonen J, et al. 2007. Quantification of the amphetamine content in seized street samples by Raman spectroscopy. J Forensic Sci. 52(1):88–92.

66. Maquelin K, Kirschner C, Choo-Smith LP, Van Den Braak N, Endtz HP, Naumann D, et al. 2002. Identification of medically relevant microorganisms by vibrational spectroscopy. J Microbiol Methods. 51(3):255–71.

