## Supplementary Information for "Raman spectroscopy reveals growth phase-dependent molecular differences in bacterial membrane vesicles"

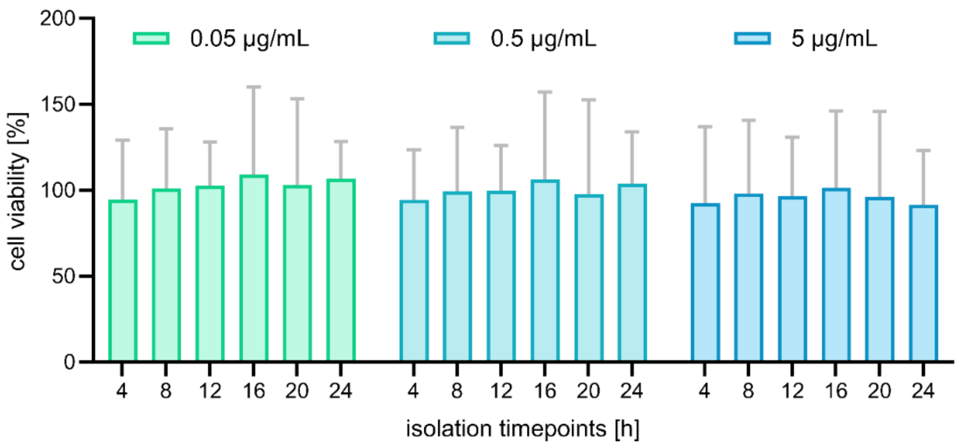


**Fig. S1**: MTT assay performed on M0 macrophages with OMVs isolated after 4, 8, 12, 16, 20, and 24 h in concentrations 0.05, 0.5, and 5 µg/mL (n = 3).

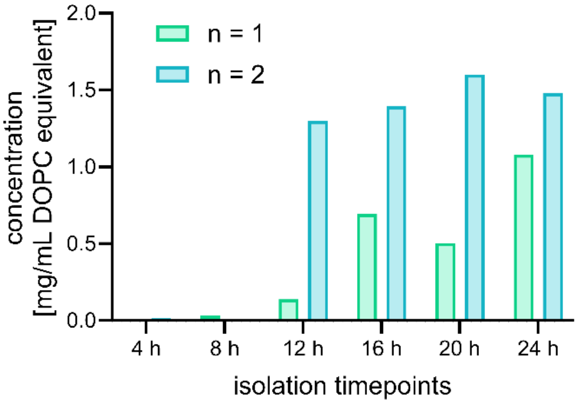


**Fig. S2**: Biochemical lipid assessment by sulpho-phospho vanillin (SPV) assay of OMVs isolated after 4, 8, 12, 16, 20, and 24 h (n = 2). Assay performed according to McMahon et al.

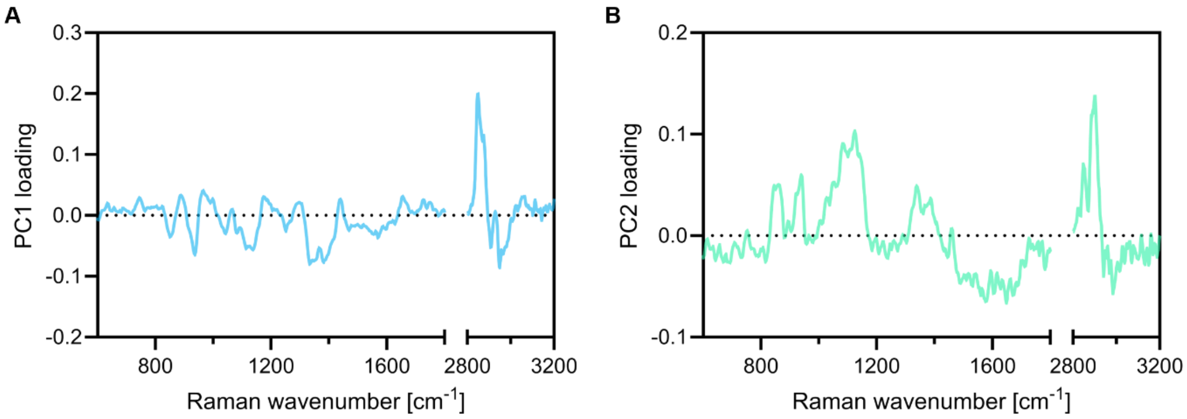


**Fig. S3**: Loading spectra for PC1 (**A**) and PC2 (**B**) used in the Principal Component Analysis in fig. 4B.

### References:

McMahon A, Lu H, Butovich IA. The spectrophotometric sulfo-phospho-vanillin assessment of total lipids in human meibomian gland secretions. Lipids, May;48(5):513-25 (2015). doi: 10.1007/s11745-013-3755-9

**Tab. S1**: Statistical evaluation by one-way ANOVA and Tukey's multiple comparison test of protein and LPS quantification shown in fig. 1B and 1C.

**One-way ANOVA and Tukey's multiple comparison test**

| Compared groups | Protein amounts (no SDS) | | Protein amounts (with SDS) | | LPS amounts | |
|---|---|---|---|---|---|---|
| | Summary | p-value | Summary | p-value | Summary | p-value |
| ANOVA summary | **** | <0.0001 | **** | <0.0001 | ns | 0.8746 |
| 4 vs. 8 | ns | 0.9960 | ns | >0.9999 | ns | 0.9992 |
| 4 vs. 12 | ns | 0.3346 | ns | 0.7736 | ns | >0.9999 |
| 4 vs. 16 | ** | 0.0022 | ns | 0.2311 | ns | >0.9999 |
| 4 vs. 20 | **** | <0.0001 | ** | 0.0014 | ns | 0.9916 |
| 4 vs. 24 | **** | <0.0001 | ** | 0.0017 | ns | 0.9768 |
| 8 vs. 12 | ns | 0.9278 | ns | 0.8511 | ns | >0.9999 |
| 8 vs. 16 | * | 0.0388 | ns | 0.3030 | ns | 0.9925 |
| 8 vs. 20 | *** | 0.0003 | ** | 0.0023 | ns | 0.9371 |
| 8 vs. 24 | **** | <0.0001 | ** | 0.0027 | ns | 0.8901 |
| 12 vs. 16 | ns | 0.6408 | ns | 0.9989 | ns | 0.9985 |
| 12 vs. 20 | * | 0.0282 | ns | 0.2168 | ns | 0.9712 |
| 12 vs. 24 | *** | 0.0008 | ns | 0.2377 | ns | 0.9405 |
| 16 vs. 20 | ns | 0.9501 | ns | 0.7542 | ns | 0.9990 |
| 16 vs. 24 | ns | 0.3302 | ns | 0.7820 | ns | 0.9951 |
| 20 vs. 24 | ns | 0.9887 | ns | >0.9999 | ns | >0.9999 |

**Tab. S2**: Statistical evaluation by one-way ANOVA and Tukey's multiple comparison test of AFM height (fig. 2B), calculated diameter (fig. 2C), hydrodynamic diameter (fig. 2D), PDI (fig. 2E), and zeta potential (fig. 2F).

**One-way ANOVA and Tukey's multiple comparison test**

| Compared groups | **AFM height** | | **Calculated diameter** | | **Hydrodyn. diameter** | | **PDI** | | **Zeta potential** | |
|---|---|---|---|---|---|---|---|---|---|---|
| | Sum. | p | Sum. | p | Sum. | p | Sum. | p | Sum. | p |
| ANOVA summary | ** | 0.0086 | ns | 0.9527 | ns | 0.2616 | *** | 0.0001 | **** | <0.0001 |
| 4 vs. 8 | ns | 0.1295 | ns | >0.9999 | ns | 0.9998 | ns | 0.8990 | ns | 0.1680 |
| 4 vs. 12 | ns | 0.1419 | ns | >0.9999 | ns | 0.8478 | ns | 0.5672 | ns | 0.0549 |
| 4 vs. 16 | ns | 0.7743 | ns | 0.9992 | ns | 0.9606 | ns | 0.0555 | *** | 0.0005 |
| 4 vs. 20 | ns | 0.1818 | ns | 0.9506 | ns | 0.9898 | *** | 0.0008 | *** | 0.0007 |
| 4 vs. 24 | ** | 0.0041 | ns | 0.9890 | ns | 0.9198 | ** | 0.0011 | **** | <0.0001 |
| 8 vs. 12 | ns | >0.9999 | ns | >0.9999 | ns | 0.9444 | ns | 0.9905 | ns | 0.9964 |
| 8 vs. 16 | ns | 0.8515 | ns | >0.9999 | ns | 0.8790 | ns | 0.4453 | ns | 0.3329 |
| 8 vs. 20 | ns | >0.9999 | ns | 0.9810 | ns | 0.9496 | * | 0.0224 | ns | 0.3914 |
| 8 vs. 24 | ns | 0.7570 | ns | 0.9977 | ns | 0.8054 | * | 0.0285 | *** | 0.0006 |
| 12 vs. 16 | ns | 0.8692 | ns | >0.9999 | ns | 0.3509 | ns | 0.8151 | ns | 0.6347 |
| 12 vs. 20 | ns | >0.9999 | ns | 0.9745 | ns | 0.4824 | ns | 0.1039 | ns | 0.6999 |
| 12 vs. 24 | ns | 0.7354 | ns | 0.9963 | ns | 0.2691 | ns | 0.1263 | ** | 0.0029 |
| 16 vs. 20 | ns | 0.9125 | ns | 0.9948 | ns | >0.9999 | ns | 0.7323 | ns | >0.9999 |
| 16 vs. 24 | ns | 0.1497 | ns | 0.9998 | ns | >0.9999 | ns | 0.7820 | ns | 0.1857 |
| 20 vs. 24 | ns | 0.6709 | ns | 0.9998 | ns | 0.9990 | ns | >0.9999 | ns | 0.1501 |

**Tab. S3**: Statistical evaluation by one-way ANOVA and Tukey's multiple comparison test of Raman peak ratios shown in fig. 4D-G.

| | One-way ANOVA and Tukey's multiple comparison test | | | | | | | |
|---|---|---|---|---|---|---|---|---|
| | **Protein-to-lipid ratio** (1645 $cm^{-1}$ / 1302 $cm^{-1}$) | | **Protein secondary structure** (1279 $cm^{-1}$ / 1235 $cm^{-1}$) | | **Lipid saturation** (2865 $cm^{-1}$ / 2935 $cm^{-1}$) | | **Carbohydrate levels** (1338 $cm^{-1}$ / 2935 $cm^{-1}$) | |
| Compared groups | Sum. | p | Sum. | p | Sum. | p | Sum. | p |
| ANOVA summary | **** | <0.0001 | **** | <0.0001 | **** | <0.0001 | **** | <0.0001 |
| 4 vs. 8 | **** | <0.0001 | * | 0.0269 | **** | <0.0001 | ns | 0.1713 |
| 4 vs. 12 | **** | <0.0001 | **** | <0.0001 | **** | <0.0001 | ** | 0.0028 |
| 4 vs. 16 | **** | <0.0001 | **** | <0.0001 | **** | <0.0001 | **** | <0.0001 |
| 4 vs. 20 | **** | <0.0001 | **** | <0.0001 | **** | <0.0001 | **** | <0.0001 |
| 4 vs. 24 | **** | <0.0001 | **** | <0.0001 | **** | <0.0001 | **** | <0.0001 |
| 8 vs. 12 | *** | 0.0001 | **** | <0.0001 | **** | <0.0001 | **** | <0.0001 |
| 8 vs. 16 | **** | <0.0001 | **** | <0.0001 | **** | <0.0001 | **** | <0.0001 |
| 8 vs. 20 | ns | 0.3365 | **** | <0.0001 | **** | <0.0001 | **** | <0.0001 |
| 8 vs. 24 | ns | 0.9900 | **** | <0.0001 | **** | <0.0001 | **** | <0.0001 |
| 12 vs. 16 | ns | 0.1574 | **** | <0.0001 | **** | <0.0001 | **** | <0.0001 |
| 12 vs. 20 | **** | <0.0001 | **** | <0.0001 | **** | <0.0001 | **** | <0.0001 |
| 12 vs. 24 | **** | <0.0001 | **** | <0.0001 | **** | <0.0001 | **** | <0.0001 |
| 16 vs. 20 | **** | <0.0001 | ns | 0.1720 | **** | <0.0001 | **** | <0.0001 |
| 16 vs. 24 | **** | <0.0001 | **** | <0.0001 | * | 0.0474 | ns | 0.0752 |
| 20 vs. 24 | ns | 0.7295 | ns | 0.0783 | **** | <0.0001 | ** | 0.0013 |

**Tab. S4**: Statistical evaluation by one-way ANOVA and Tukey's multiple comparison test of ELISA data on pro-inflammatory cytokines IL-1β (fig. 5A), IL-6 (fig. 5B), and TNFα (fig. 5C).

**One-way ANOVA and Tukey's multiple comparison test**

| Compared groups | **IL-1β** | | **IL-6** | | **TNF-α** | |
|---|---|---|---|---|---|---|
| | Summary | p-value | Summary | p-value | Summary | p-value |
| ANOVA summary | **** | <0.0001 | **** | <0.0001 | **** | <0.0001 |
| 4 vs. 8 | ns | 0.9299 | ns | >0.9999 | ns | 0.3261 |
| 4 vs. 12 | *** | 0.0009 | ns | >0.9999 | ns | 0.9929 |
| 4 vs. 16 | ** | 0.0024 | ns | >0.9999 | ns | 0.0799 |
| 4 vs. 20 | **** | <0.0001 | ns | >0.9999 | ns | 0.0914 |
| 4 vs. 24 | **** | <0.0001 | ns | >0.9999 | * | 0.0294 |
| 4 vs. (-) | **** | <0.0001 | *** | 0.0001 | **** | <0.0001 |
| 4 vs. (+) | *** | 0.0003 | ns | 0.9994 | ns | >0.9999 |
| 8 vs. 12 | ** | 0.0086 | ns | >0.9999 | ns | 0.7510 |
| 8 vs. 16 | * | 0.0236 | ns | >0.9999 | ns | 0.9861 |
| 8 vs. 20 | **** | <0.0001 | ns | >0.9999 | ns | 0.9918 |
| 8 vs. 24 | **** | <0.0001 | ns | >0.9999 | ns | 0.8483 |
| 8 vs. (-) | **** | <0.0001 | *** | 0.0001 | **** | <0.0001 |
| 8 vs. (+) | ** | 0.0028 | ns | 0.9985 | ns | 0.2445 |
| 12 vs. 16 | ns | 0.9994 | ns | >0.9999 | ns | 0.2850 |
| 12 vs. 20 | * | 0.0325 | ns | >0.9999 | ns | 0.3176 |
| 12 vs. 24 | ns | 0.0934 | ns | >0.9999 | ns | 0.1199 |
| 12 vs. (-) | *** | 0.0001 | *** | 0.0001 | **** | <0.0001 |
| 12 vs. (+) | ns | 0.9987 | ns | 0.9994 | ns | 0.9742 |
| 16 vs. 20 | * | 0.0119 | ns | 0.9978 | ns | >0.9999 |
| 16 vs. 24 | * | 0.0358 | ns | 0.9998 | ns | 0.9992 |
| 16 vs. (-) | **** | <0.0001 | **** | <0.0001 | **** | <0.0001 |
| 16 vs. (+) | ns | 0.9481 | ns | 0.9865 | ns | 0.0558 |
| 20 vs. 24 | ns | 0.9988 | ns | >0.9999 | ns | 0.9982 |
| 20 vs. (-) | ns | 0.1438 | *** | 0.0002 | **** | <0.0001 |
| 20 vs. (+) | ns | 0.0953 | ns | >0.9999 | ns | 0.0641 |
| 24 vs. (-) | ns | 0.0518 | *** | 0.0002 | **** | <0.0001 |
| 24 vs. (+) | ns | 0.2466 | ns | 0.9998 | * | 0.0202 |
| (-) vs. (+) | *** | 0.0004 | *** | 0.0004 | **** | <0.0001 |
